# Psilocybin ameliorates neuropathic pain-like behaviour in mice and facilitates gabapentin-mediated analgesia

**DOI:** 10.1101/2025.09.15.676273

**Authors:** Tatum Askey, Daniel Allen-Ross, Daniil Luzyanin, Reena Lasrado, Gary Gilmour, Stephen P Hunt, Francesco Tamagnini, Maqsood Ahmed, Gary J Stephens, Maria Maiarú

## Abstract

Chronic pain states are challenging to control with current drug therapies. Here, we demonstrate that a single dose of psilocybin can produce a sustained anti-nociceptive effect in a model of chronic neuropathic pain in male and female mice. Psilocybin anti-nociceptive effects were mediated by 5-HT_2A_ receptors, although additional mechanisms might also be involved. Furthermore, a single dose of psilocybin caused a significant increase in the anti-nociceptive potential of gabapentin, a widely used treatment for neuropathic pain consistent with the establishment of longer lasting changes in network processing. Overall, these findings present the first preclinical evidence that psilocybin could be a valuable approach for treating chronic pain from nerve injury and serve as a new therapeutic addition for pain management.

## Main Text

Chronic pain affects millions of people worldwide and presents a huge social and economic burden. Long lasting pain negatively affects the quality of life of patients^1^ and is associated with significant unmet clinical need. Management of chronic pain is difficult and clinically available drugs are often poorly tolerated and/or potentially addictive^2,3^. People with chronic pain often develop affective comorbidities, such as depression and anxiety^4^ that are frequently associated with worsening clinical outcomes^5^.

Psilocybin is a classic psychedelic drug that typically causes a profound alteration of perception and mood^6^. The active metabolite of psilocybin is psilocin, which is known to bind to multiple serotoninergic receptors, of which the 5-hydroxytryptamine (5-HT) 2A receptor (5-HT_2A_R) is necessary for the induction of the psychedelic effect^7^. There has been a resurgence of interest in the clinical potential of psilocybin for mental health conditions such as major depression disorders^8^, diseases that are typically co-morbid with chronic pain states. Positive treatment outcomes following a single treatment with psylocibin have been associated with the long-term modulation of intrinsic brain networks and the resetting of aberrant connectivity that reinforced negative patterns of behaviour^9^.

There are also some early indications that psilocybin may alleviate intractable phantom limb pain^10^, migraine and associated pain^11^ in humans and reduce mechanical hypersensitivity in an inflammatory pain model in male and female rats^12^. Investigating neuronal networks that might sustain chronic pain states remains at an early stage^13^, but maladaptive changes to functional and structural connectivity have been demonstrated to precede the onset of chronic subacute lower back pain in human patients^14^. This led us to consider the value of psilocybin as a potential treatment for ‘unlocking’ neuronal networks that support chronic neuropathic pain^15^ and to evaluate whether 5-HT_2A_R activation is necessary for this response. Moreover, we also examined the hypothesis that psilocybin could positively affect the anti-nociceptive response of an analgesic established as a treatment option for neuropathic pain.

## Results

### Psilocybin shows anti-nociceptive effects in male mice after injury

We induced a chronic pain-like status in adult C57BL/6J male and female mice. The spared nerve injury (SNI) mouse model is a preclinical model of neuropathic pain involving partial transection of the peripheral nerves innervating the hind paw^16^. The SNI model is considered one of the best experimental models for studying neuropathic pain induced by nerve injury due to its relevance to human pain conditions and its ability to reproduce several key aspects of neuropathic pain^17^. This model mimics the type of nerve injury commonly seen in human neuropathic pain conditions, such as those resulting from traumatic injury, surgery, or diseases such as diabetes. The model selectively injures the sciatic nerve, creating a partial nerve injury rather than a complete transection, which preserves some nerve fibers while damaging others. This partial injury is representative of many clinical neuropathic pain conditions where nerves are not entirely severed but are damaged to varying degrees. Moreover, the pain behaviours that develop as a result of the injury, such as mechanical hypersensitivity (allodynia) and thermal hypersensitivity (hyperalgesia), are similar to those observed in humans with neuropathic pain, making it an excellent model for studying pain and responses to analgesics.

The head-twitch response (HTR) is considered a rodent behavioural proxy of the human response to a psychedelic experience^18^ and psilocybin administration has been shown to induce HTR in mice^19^, with a marked increase in this behaviour observed at a dose of 1 mg/kg. In a dose-response curve, the HTR seen after a 1 mg/kg dose was comparable to that induced by a higher dose of psilocybin^19^; based on this, we selected doses of 1 mg/kg and 0.3 mg/kg of psilocybin in the SNI model to evaluate its effects across a range of different behavioural tests that measure both reflexive and affective responses to mechanical and thermal stimulation of the hind paw.

The experimental design is summarised in **Figure 1 a**. Male mice underwent SNI surgery and, when static mechanical hypersensitivity was fully developed (day 12), they received a single intraperitoneal (i.p.) injection of psilocybin (0.3 mg/kg) or saline control (**Fig 1 a**). An increase in HTR was observed in the psilocybin treatment groups compared to saline control (**Fig 1 b**) confirming central nervous system exposure to the drug. Mechanical hypersensitivity was reduced (Maximum Possible Effect, MPE=18%) in mice treated with psilocybin with the effect lasting up to day 28 after injection (test day 40) (**Fig 1 c**). Moreover, preliminary observations were that affective behaviours that characterise dynamic mechanical hypersensitivity to light brush stroke after psilocybin treatment were reduced (**Fig 1 d**), but that no effect was observed on the licking/biting response to a cold stimulus (**Fig 1 e**). The single dose of psilocybin (0.3 mg/kg) had no effect on locomotor performance in SNI mice (**Fig S1**), consistent with psilocybin not causing unwanted motor function deficits. Taken together, we observed a number of anti-nociceptive effect of psilocybin in the SNI model, consistent with the potential alleviative effects in the human pain experience.

**Fig. 1:**
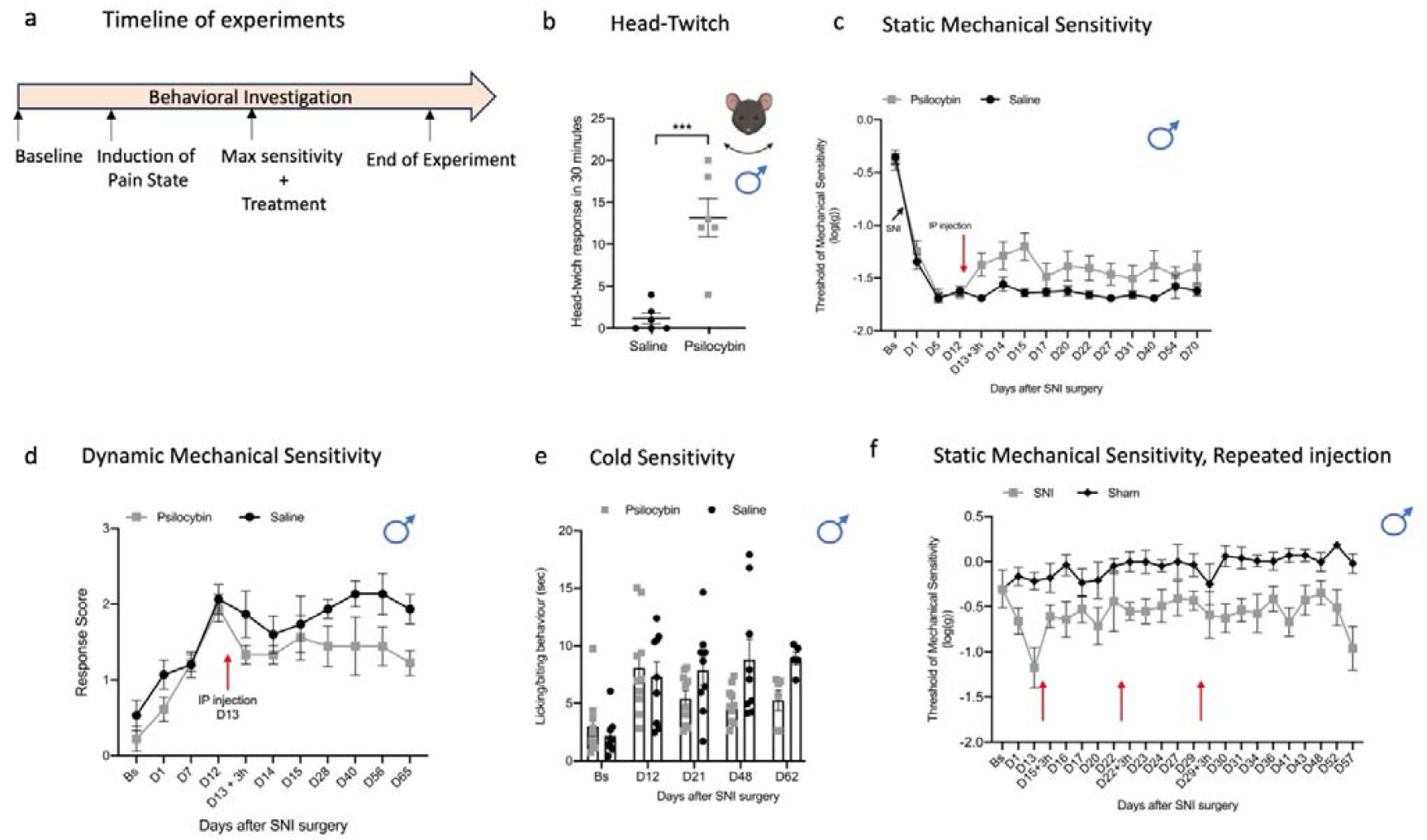
Psilocybin (0.3 mg/kg) attenuates pain-like behaviour in male mice after peripheral nerve injury. **a**, Schematic of timeline of the experiments (Bs, Baseline). **b**, Head-twitch response after injection of psilocybin (0.3 mg/kg) (n=6) or saline control (n=6). P=0.001, unpaired independent sample t-test. **c**, Static mechanical threshold of mice assessed using calibrated von Frey filaments before (Bs, baseline) and after SNI surgery (D1 to D40) (n=10/9, two-way, repeated measured with mixed models ANOVA, factor TREATMENT D13+3h to D54: F= 10.7, P=0.004). **d**, Brush-evoked dynamic hypersensitivity before (Bs) and after SNI surgery (n=6/5, two-way, repeated measured with mixed models ANOVA, factor TREATMENT D13 to D65: F= 7.2, P = 0.025). **e**, Cold allodynia assessed using acetone drop applied to the hindpaw ipsilateral to the injury (left) before (Bs) and after SNI surgery (n= 10/9, two-way, repeated measured with mixed models ANOVA, factor TREATMENT D21 to D62: F= 2.8, P = 0.13). **f**, Static mechanical threshold of mice assessed using calibrated von Frey filaments before (BS, baseline) and after SNI or SHAM surgery. On day 15, 22 and 29 after SNI, all mice received an IP injection of psilocybin (0.3 mg/kg) red arrows (n=5/5, two-way, repeated measured with mixed models ANOVA, factor TREATMENT D15+3h to D57: F= 13.7, P = 0.006). Data are expressed as mean ± SEM throughout. Red arrows represent psilocybin injection. SNI, Spared Nerve Injury; IP, Intra-peritoneal.

We next tested whether repeated injection of psilocybin (0.3 mg/kg) could amplify the anti-nociceptive effects observed with a single dose. Psilocybin was injected every 7 days for 3 weeks in male mice that had undergone SNI or sham surgery (**Fig 1 f**). A reduction of mechanical hypersensitivity was observed in SNI mice. Repeated injections of psilocybin (0.3 mg/kg) substantially prolonged and amplified the antinociceptive effects for several weeks (**Fig 1 f**) in comparison to a single dose (**Fig 1 c**) (%MPE= 62.6 in the same group of mice (SNI) before and after psilocybin injection). No changes in mechanical threshold were observed in sham mice (**Fig 1 f**).

### Higher-dose psilocybin improves pain and enhances gabapentin effects

In the next series of experiments, a single higher dose of psilocybin (1 mg/kg i.p.) induced a robust HTR in male and female mice (**Fig 2 a**). At this dose, psilocybin-induced reduction in mechanical hypersensitivity (MPE=25.5%) was comparable to the lower dose (MPE=18% 0.3 mg/kg) in male mice and similarly persisted for several days after injury (**Fig 2 b**). The single dose of psilocybin (1 mg/kg) had no adverse effect on locomotor performance in SNI mice (**Fig 2 c**). In male mice, preliminary data also indicate that psilocybin (1 mg/kg) was also able to reduce hypersensitivity to light brush that develops after peripheral nerve injury (**Fig 2 d**). To assess cold sensitivity, we used the thermal place preference test (TPP)^21^ and recorded the total amount of time mice spent on the cold plate before (Bs) and after nerve injury (day 1 to day 30) (**Fig 2 e**). A overall trend toward an increase in the time spent on the cold plate was observed (P=0.085), compared to saline-injected mice which reached significance by day 30 (Bonferroni corrections P=0.02) (**Fig 2 e**) (psilocybin, 65.1 ± 23.4 s; saline, 18.9 s ±3.6 s) Psilocybin (1 mg/kg) also had no adverse effect on locomotor performance in naïve male mice (**Fig S2**). Next, we analysed faecal output as a measure of stress in mice^22^ after SNI for psilocybin (1 mg/kg) vs saline controls. Psilocybin treatment reduced faecal boli output in mice after SNI surgery (**Fig S3**), this was also associated with increased body weight (**Fig S3**). These data are consistent with the hypothesis that psilocybin may reduce the stress that follows injury, this aspect warrants further investigation.

**Fig 2:**
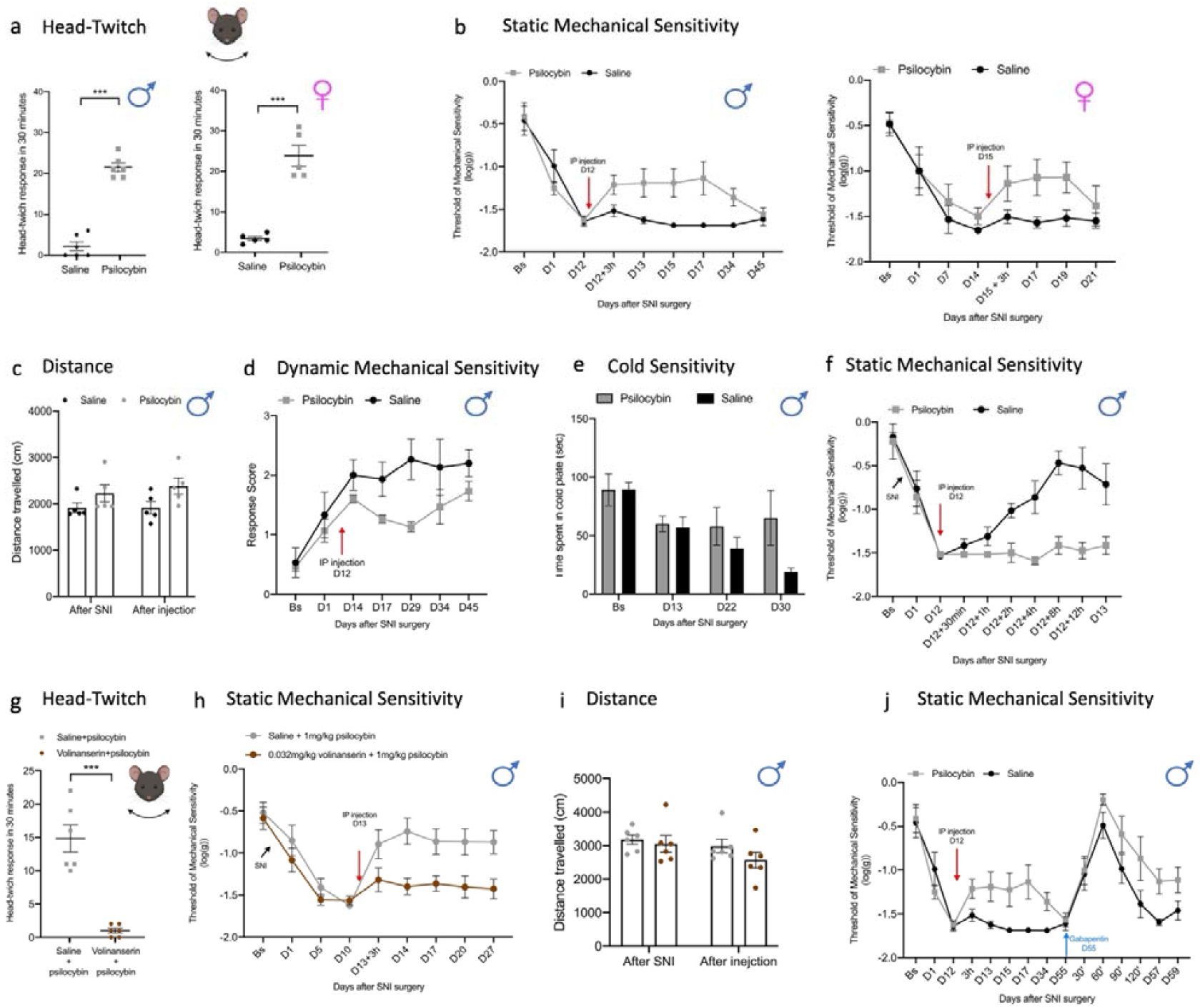
Psilocybin (1 mg/kg) improves hypersensitivity in male and female mice and potentiates the effect of gabapentin. **a**, Head-twitch response in male and female mice after injection of psilocybin (1 mg/kg) (male, n=6; female, n=5) or saline control (male, n=6; female, n=5); P < 0.001 unpaired independent sample t-test. **b**, Static mechanical threshold of mice assessed using calibrated von Frey filaments before (Bs, baseline) and after SNI surgery. At maximum sensitivity day 12 (male) or on day 15 (female), mice received an IP injection of 1 mg/kg psilocybin or saline control (male, n=8/8, two-way repeated measured with mixed models ANOVA, factor TREATMENT 3h to D45: F= 10.05, P=0.007; female, n=8/8, two-way repeated measured with mixed models ANOVA, factor TREATMENT D15+3h to D21: F= 12.1, P=0.005) (on D21 psilocybin n=5). **c**, Locomotor activity (distance travelled) after SNI surgery and subsequent injection of saline or psilocybin. Bar graph displays the total distance travelled (cm) by animals following SNI (“After SNI”) and after acute injection (“After injection”) of either saline (black circles) or psilocybin (grey circles). Data are presented as mean ± SEM, with individual data points shown (n=5/5). **d**, Brush-evoked dynamic hypersensitivity before (Bs) and after SNI surgery (n=5/5, two-way repeated measured with mixed models ANOVA, factor TREATMENT D14 to D45: F= 6.5, P=0.034). **e**, Cold allodynia assessed using Thermal Preference Test before (Bs) and after SNI surgery (n= 5/5, two-way repeated measured with mixed models ANOVA, factor TREATMENT D13 to D30: F= 3.9, P=0.085, with Bonferroni corrections D22, P=0.09, D30, P=0.02). **f**, acute effect of psilocybin on static mechanical sensitivity assessed using calibrated von Frey filaments. At maximum sensitivity day 12 mice received an IP injection of 1 mg/kg psilocybin or saline control. Behavioural measures taken at several time points after injections (30 min, 1h, 2h, 4h, 8, 12h and 24h). (n=6/6, two-way repeated measured with mixed models ANOVA, factor TREATMENT 30min to D13: F= 18.7, P=0.002; with Bonferroni corrections 1h, P=0.085, 2h, P=0.005, 4h, P=0.004, 8h, P<0.001, 12h, P=0.004, 24h, P=0.022). **g**, Head-twitch response in male mice after injection of psilocybin (1 mg/kg) and volinanserin (0.032 mg/kg) (n=6/6, P < 0.001 unpaired independent sample t-test). **h**, Static mechanical threshold of mice before (BS, Baseline) and after SNI surgery (D1 to D27). On day13 mice received an injection of 0.032mg/kg of volinanserin or saline control; 30 minutes later all mice received an IP injection of 1 mg/kg of psilocybin. (n=12/12, two-repeated measured with mixed models ANOVA, factor TREATMENT D13+3h to D27: F= 11.9, P=0.002). **i**, Locomotor activity (distance travelled) after SNI surgery and subsequent injection of saline or psilocybin. Bar graph displays the total distance travelled (cm) by animals following SNI (“After SNI”) and after acute injection (“After injection”) of either saline + psilocybin (grey circles) or volinanserin + psilocybin (brown circles). Data are presented as mean ± SEM, with individual data points shown (n=6/6). **j**, Psilocybin (1 mg/kg) or saline control were injected intraperitoneal (i.p.) 12 days after SNI surgery (red arrows). Gabapentin (50mg/kg) was injected i.p. 55 days after SNI surgery (blue arrow). (n=8/8, two-way repeated measured with mixed models ANOVA, factor TREATMENT 60’ to D59: F_1,14_ = 9.59, P=0.008). Data are expressed as mean ± SEM throughout. Red arrows represent psilocybin injection. Blue arrow represents injection gabapentin injection. SNI, Spared Nerve Injury; IP, Intra-peritoneal.

To further characterise the temporal profile of psilocybin’s effects on mechanical sensitivity, we performed a detailed time-course from 30 minutes to 24 hours post-injection (**Fig 2 f**). No significant effect of psilocybin was observed at 30 minutes and 1-hour post-injection; however, psilocybin-treated mice showed robust and sustained reductions in mechanical sensitivity from 2 hours onwards, which persisted throughout the 24-hour assessment period (MPE=46%). By contrast, saline-treated mice did not show changes in mechanical sensitivity over the same timeframe.

We next tested whether 5-HT_2A_R activation is required for the anti-nociceptive effect of psilocybin. In mice, psilocybin-induced HTR is mediated by the 5-HT_2A_Rs^18^. We therefore pre-injected male mice with the 5-HT_2A_R antagonist volinanserin (0.032mg/kg i.p.), shown to be effective in previous mouse behavioural studies^23^, followed 30 mins later by psilocybin (1 mg/kg i.p.). Control mice received saline prior to psilocybin. Volinanserin pretreatment blocked the HTR in mice (**Fig 2 g**) and abolished its full anti-nociceptive effect on mechanical hypersensitivity (**Fig 2 h**), but had no adverse effect on locomotory functions (**Fig 2 i**). Volinanserin alone has no effect on mechanical hypersensitivity (**Fig S4**).

We also assessed the effect of psilocybin and volinanserin treatments on spontaneous behaviour in naïve mice (**Fig S5**). No differences were observed in total distance travelled across groups, supporting that neither psilocybin nor volinanserin impaired locomotor functions. Unsupported rearing and grooming behaviours were reduced by psilocybin 10 minutes post-injections, but these changes were absent 3 days later. Volinanserin did not cause any additional changes in grooming and unsupported rearing in naïve mice, consistent with lack of effect on locomotory activity; these data also suggest that these behaviours may be mediated by a 5HT_2A_R independent mechanism^24^.

The neural mechanisms underlying the effects of psilocybin are not fully understood but are thought to involve the modulation of normal patterns of communication between different areas of the brain. Altered brain functional connectivity in chronic pain patients and mice have been observed^25,26^ and the analgesic effect of psychedelic drugs could be due to their capacity to drive neuroplasticity and reset aberrant connections that support chronic pain^27^. Given that the effects of a single dose of psilocybin can last for many months in both people with depression^28^ and control groups, we hypothesised that psilocybin might be able to influence pain processing networks in mice beyond the period when alteration in pain behaviours were seen. We therefore investigated the effect of psilocybin on response to gabapentin. Mice received a single i.p. injection of gabapentin (50 mg/kg i.p.) at day 55 after surgery when the anti-nociceptive effect of psilocybin was no longer measurable (**Fig 2 j**). In mice treated with psilocybin (1 mg/kg i.p.), a dramatic prolonged anti-nociceptive effect of gabapentin from 2h to 96h was observed compared to mice treated with saline vehicle (**Fig 2 j**).

## Discussion

Developing safer, effective treatments for chronic pain has proven challenging. Here, we demonstrate that psilocybin can reduce neuropathic pain-like behaviour in male mice for up to 30 days, that this reduction in pain behaviour can be amplified by repeated treatment with low dose psilocybin and finally that a later time point psilocybin can potentiate the effect of gabapentin, a standard treatment for neuropathic pain in humans. Recent research has reported no general differences between males and females in response to psilocybin in a rat model of inflammatory pain^12^ and here we show that psilocybin also reduces mechanical sensitivity in female mice in the SNI model of neuropathic pain. A single dose of psilocybin also had a persistent effect in female mice, but a fuller characterisation of psilocybin anti-nociceptive effects in female animals is warranted to determine if sex differences, such as the duration of effect on mechanical sensitivity we report here, extend to other measures.

Notably, a recent preprint reported that psilocybin at doses as high as 10 mg/kg lacked analgesic effects in mixed male and female cohorts in several mouse pain models^29^; this is in contrast to reported anti-nociceptive effects in models of chemotherapy-induced peripheral neuropathy (CIPN)^30^ and inflammatory pain^12^ (reviewed in^15^) and in the SNI model here. Our experimental design facilitated detection of neuroplasticity-driven pain reduction that might not be captured in acute paradigms and include an initial consideration of sex differences.

Human studies have confirmed that 5-HT_2A_R activation is responsible for the profound changes in perception and consciousness of psilocybin, but the role of 5-HT_2A_Rs in mediating clinical efficacy remains to be determined fully. Rodents studies have established that 5-HT_2A_R activation is necessary for HTR, and our data demonstrates that volinanserin, a selective 5-HT_2A_R antagonist, can prevent the full anti-nociceptive effect of psilocybin. By contrast, one major earlier behavioural study reported that ketanserin, a 5-HT_2A_/_2C_R antagonist, did not influence psilocybin’s antidepressant-like behaviours, suggesting that the receptor mechanisms underlying different therapeutic effect of psilocybin may be distinct^31,32^. Other studies using ketanserin indicate an, at least partial, involvement of 5HT_2A_Rs in schizophrenia-like psychosis^33^ and cognitive functions^34^. Psilocin also has appreciable affinity at other 5HTRs, including 5-HT_1A_R^35^. Recent research using volinanserin has shown complete blockade of psilocybin’s antinociceptive effects in chemotherapy-induced peripheral neuropathy (CIPN) models^30^. In these studies, volinanserin pretreatment (0.05 mg/kg) completely blocked the anti-nociceptive effects of both psilocybin and DOI, indicating that 5-HT_2A_R activation is necessary for psilocybin’s analgesic properties in neuropathic pain models. These findings support the analgesic properties of psilocybin require 5-HT_2A_R activation.

Our data show that volinanserin does not fully block psilocybin anti-nociceptive effects in SNI model; these data suggest that other mechanisms may also contribute. In this regard, recent research has suggested that psilocybin may directly bind to the neuronal receptor tyrosine kinase B (TrkB) receptor and allosterically potentiate BDNF (brain derived neurotropic factor) signalling, independently of 5HT_2A_R activation^36,37^. Neurons in both the frontal cortex and dorsal horn of the spinal cord have been shown to responds to psilocybin by augmentation of BDNF signalling^36^. Psilocybin induces a rapid and persistent growth of dendritic spines of medial prefrontal cortex neurons^19^ which may contribute to the increased excitability of neurons and a reduction in the chronic pain behaviour reported^25^. Future investigations are therefore required to establish whether the anti-nociceptive effect of psilocybin could also be linked to the action of BDNF released in the cortex and dorsal horn.

Gabapentin is a drug widely used in clinical practice to treat neuropathic pain; however, not all the people with neuropathic pain achieve adequate pain relief with gabapentin^38^; moreover, gabapentin use is also associated with side effects^38^ and a risk of addiction^39^. A striking finding was the dramatic and sustained enhancement of gabapentin’s anti-nociceptive efficacy when administered weeks after psilocybin treatment, at a time when psilocybin’s direct ani-nociceptive effects were no longer detectable. One potential explanation is that psilocybin may induce persistent neuroplastic changes^19,40^ that alter pain processing networks; this may create a neurobiological environment more conducive to gabapentin’s therapeutic actions. Several neurobiological mechanisms could explain psilocybin’s ability to enhance gabapentin analgesic efficacy weeks after administrations. For example, BDNF expression in the rostral ventromedial medulla (RVM) has been shown to be essential for morphine’s analgesic effects and can potentiate morphine efficacy at otherwise ineffective doses^37^. The synergism observed here between psilocybin and gabapentin strongly justifies investigating whether similar enhancement occurs with other drugs used to target chronic neuropathic pain such as morphine, amitriptyline and duloxetine^41^.

Together, these data provide the first preclinical demonstration that psilocybin could be an effective tool for the management of chronic pain due to nerve injury across sexes and offer a new therapeutic adjunct for the control of chronic pain. The dramatic potentiation of gabapentin efficacy weeks after psilocybin administration represents a finding that warrants immediate investigation with morphine and other analgesics. A neuroplasticity-based enhancement strategy could significantly enhance chronic pain management by improving existing treatments rather than requiring entirely new drug development. The convergent evidence from 5HT_2A_R involvement and potential network connectivity changes provides a strong mechanistic foundation to address critical clinical needs in pain medicine while contributing to our understanding of psychedelic-induced therapeutic utility.

## Material and Methods

### Experimental design

This study was designed to evaluate the effect of psilocybin on pain sensitivity. In all experiments, mice were randomly assigned into treatment groups to ensure unbiased allocation. Allocation details were concealed, and cages were coded so that the experimenter performing the behavioural assessments and analyses remained fully blind to treatment conditions throughout the study. Blinding was maintained during drug preparation, testing, and statistical evaluation to minimize potential bias. The exact number of mice per group is specified in the respective figure legends.

### Animals

All experimental procedures were conducted in accordance with institutional and national guidelines for the care and use of laboratory animals, were approved by the relevant ethical review board, and are reported in compliance with the ARRIVE (Animal Research: Reporting of In Vivo Experiments) guidelines^42^. All procedures were performed in accordance with the UK Animals (Scientific Procedures) Act, 1986, and the principles of the 3Rs (Replacement, Reduction, and Refinement). Every effort was made to minimize potential suffering, and the number of animals used was reduced to the minimum required to ensure sufficient statistical power. Sample size calculations were informed by variance estimates and effect sizes obtained from our previous behavioural assays, ensuring that group sizes were adequate for reliable detection of treatment effects without unnecessary animal use.

Adult male and female C57BL/6J mice (8–12 weeks old) were obtained from Charles River Laboratories (UK) and acclimatized to the housing facility for at least one week prior to experimentation. Animals were group-housed in individually ventilated cages (maximum of five per cage) under standard laboratory conditions, with controlled ambient temperature (20□±□1 °C) and humidity (55□±□5%). A 12:12 h light– dark cycle was maintained (lights on at 07:30 a.m.), and animals had free access to food and water throughout the study. Environmental enrichment (nesting material and shelters) was provided in all cages to promote welfare.

All experimental protocols were approved by the institutional Animal Welfare and Ethical Review Body (AWERB) and were carried out under UK Home Office Project Licence PPL PP9720547.

### Mouse model of neuropathic pain: spared nerve injury (SNI)

The SNI surgery was performed as previously described^16^. Briefly, under isoflurane anaesthesia, the skin on the lateral surface of the thigh was incised, and a section made directly though the biceps femoris muscle, exposing the sciatic nerve and its three terminal branches: the sural, the common peroneal, and the tibial nerves. The common peroneal and the tibial nerves were tightly ligated with 5-0 silk suture and sectioned distal to the ligation. Great care was taken to avoid any contact with the spared sural nerve. Complete haemostasis was confirmed, and the wound was sutured.

### Drugs

Psilocybin (COMP360, a proprietary formulation of synthetic psilocybin) was provided by Compass Pathfinder (a subsidiary of Compass Pathways) and was dissolved in saline. Control mice received equal volume of saline.

Gabapentin was purchased from Sigma and was dissolved in saline.

Volinanserin was purchased from Merck and was dissolved in saline.

### Behavioural testing

#### Von Frey filament test for static mechanical sensitivity

For the assessment of mechanical sensitivity, the von Frey filament test was used as previously described^20^. Mice were placed in Plexiglas chambers, located on an elevated grid, and allowed to habituate for at least 1h. After this time, the plantar surface of the paw was stimulated with a series of calibrated von Frey monofilaments, mechanical sensitivity threshold was determined using the up-down-method^43^. The data were expressed as log of the mean of 50% pain threshold ± SEM.

#### Brush test for dynamic mechanical sensitivity

Dynamic allodynia was tested by light stroking (velocity ∼2cm/s) of the external lateral side of the injured hind paw in the direction from heel to toe using a paintbrush using a protocol adapted from Duan and colleagues^44^. Mice were placed in Plexiglas chambers located on an elevated grid, and allowed to habituate for at least 1h. Observed responses were scored: 0, no response or moving the stimulated paw; 1, single withdrawal, flick or stamp of the stimulated paw; 2, multiple withdrawals of the stimulated paw in rapid succession; 3, licking of the plantar surface or continued elevation/withdrawal of the stimulated paw. The stimulation was repeated three times at intervals of at least 3min and the average scores was obtained for each mouse.

#### Acetone test for cold sensitivity

For assessment of cold sensitivity, the acetone test was used as previously described^20^. Mice were placed in Plexiglas chamber located on an elevated grid for 1h and then a drop (∼50μl) of acetone was applied to the external lateral side of the injured hind paw. Total time licking/biting of the hid paw was recorded for 20s.

#### Rotarod test

Locomotor activity was analysed as previously described^45^. Briefly, in this study we used an accelerating rotarod apparatus with a 3cm diameter rod starting at an initial rotation of 4rpm and slowly accelerating to 40rpm over 100s. Mice were expected to walk at the speed of rod rotation to keep from falling. Mice were not tested at baseline to minimize the number of tests on the apparatus. Time taken to fall from the rod was recorded.

#### Fecal pellet output

Mice were placed in Plexiglas chamber located on an elevated grid for 1h, after this time the number of fecal pellets were counted for each mouse.

#### Head twitch response

After the drug or saline was administered, the mice were placed in individual Plexiglas chambers and movement recorded using a digital camera. The videos for each mouse were subsequently analyzed to count the number of head twitches, which are characterized by rapid, involuntary movements of the head. The number of head twitches was counted over a 30 min period.

#### Spontaneous behaviour

Spontaneous behaviour was assessed using the EthoVision XT video tracking system. Each animal was placed individually in the open-field arena, and behaviour was recorded for the duration of the test session (10min). Automated tracking was used to quantify total distance travelled (cm) as a measure of locomotor activity. In addition, ethological parameters including unsupported rearing and grooming episodes were coded and analysed, providing indices of exploratory behaviour and self-maintenance, respectively. All recordings were performed under consistent lighting and environmental conditions to minimize variability.

#### Open Field Test

The open field test (OFT) was used to analyze the effect of psilocybin on locomotor activity (measured as the total distance travelled and the frequency of entering the centre zone). Mice were tested 24 h post-injection with 1 mg/kg of psilocybin or saline vehicle. Mice were placed in the centre of a circular plastic box (33 cm × 33 cm) and allowed to move freely. The total distance travelled and the frequency of entering the centre of the arena were recorded over the next 5 min; the EthoVision Track System was used to record and analyze the behaviour.

#### Data and statistical analysis

All statistical tests were performed using the IBM SPSS Statistic Programme (version 27), and P<0.05 was considered statistically significant. Data are means ± s.e.m. and independent experiment unit (n) is animals. As previously described^20^, data of von Frey filaments test was log transformed to ensure a normal distribution^46^.

Differences in sensitivity were assessed using repeated-measures mixed-model ANOVA, with “time” specified as a within-subject factor and “treatment” as a between-subject factor. Following a significant interaction or main effect, pairwise comparisons were performed using Bonferroni correction to control for multiple testing. Where comparisons across all group means were required, a Tukey’s post-hoc test was applied.

The MPE (Maximum Possible Effect) was calculated as previously described and according to the formula:

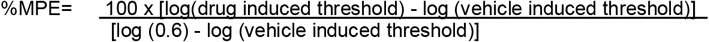

where log(0.6) is our maximum von Frey’s force applied.

## Data availability

Source data are provided with this paper and available at UoR Research Data Archive

## Contributions

TA, MA, SPH, GJS and MM conceived and designed the experiments; TA, DAR, DL and MM performed the experiments; TA and MM analysed the data; TA, MA, SPH, GJS and MM drafted the paper and MM wrote the manuscript. All authors revised and edited the manuscript.

## Supplementary figures

**Fig S1.**
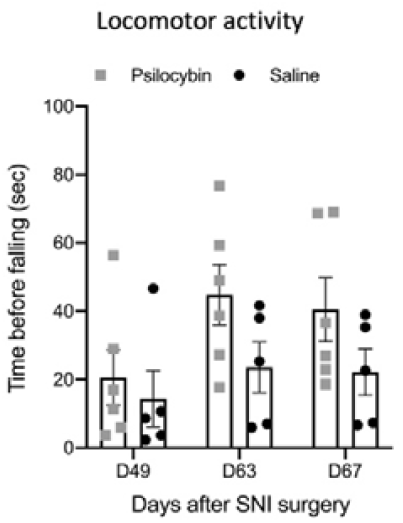
Effect of psilocybin on locomotor activity in male mice after SNI surgery. Locomotor activity measured as time before falling (in seconds) at three time points (Day 49, Day 63, and Day 67) after SNI surgery after psilocybin or saline treatment. Bars represent mean ± SEM. Individual data points are illustrated for each group at each time point.

**Fig S2.**
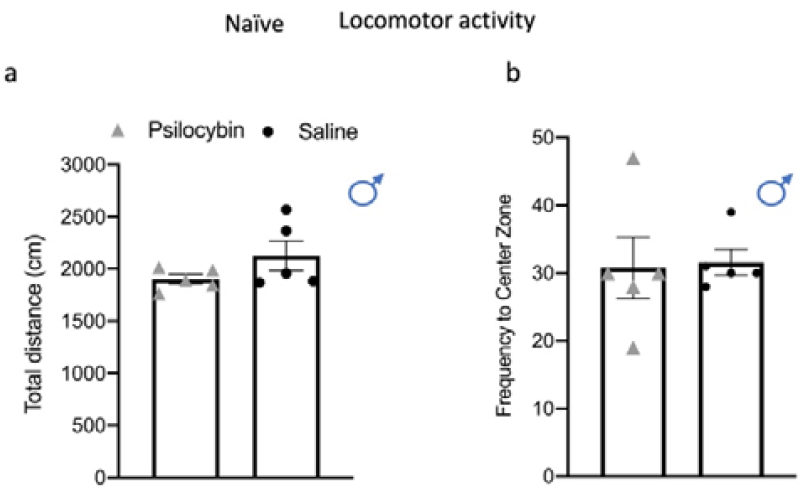
Effect of psilocybin on locomotor activity in naïve male mice. The Open Field Test was used to test locomotory activity after injection of psilocybin (1 mg/kg) or saline vehicle control in naïve male mice. The total distance travelled (**a**) and the frequency to enter the centre zone (**b**) were unaffected by psilocybin. n=5/5.

**Fig S3.**
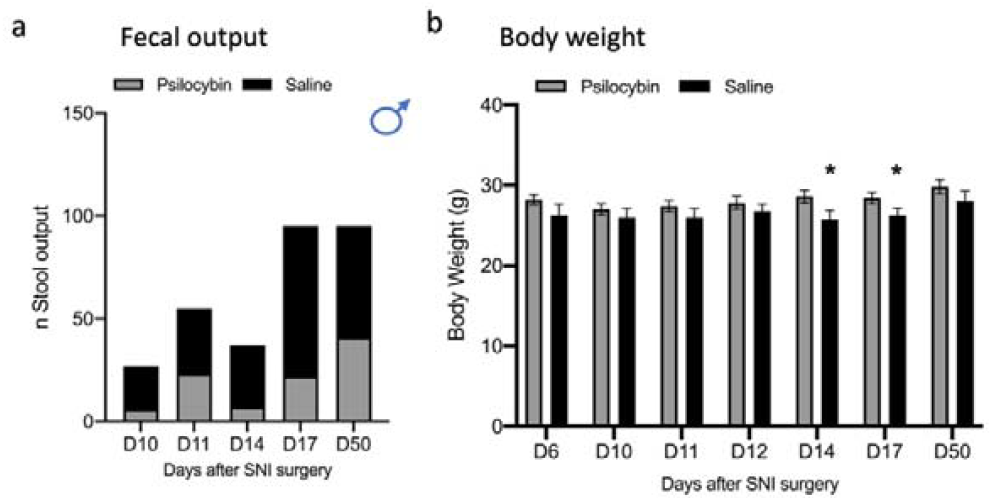
Effect of psilocybin on fecal output and body weight of male mice after peripheral nerve injury. **a**. Total stool output was measured in male mice at different time points following spared nerve injury (SNI) surgery (days 10, 11, 14, 17, and 50). Bars represent cumulative stool counts from mice treated with psilocybin (grey) or saline (black). **b**, Body weight was recorded before behavioural tests. SNI surgery was performed on D0 and psilocybin (1 mg/kg) or saline treatment was given on D12 after surgery. n=5/5. ^*^P<0.05, Student’s t-test. Data are expresses as mean ± SEM.

**Fig S4:**
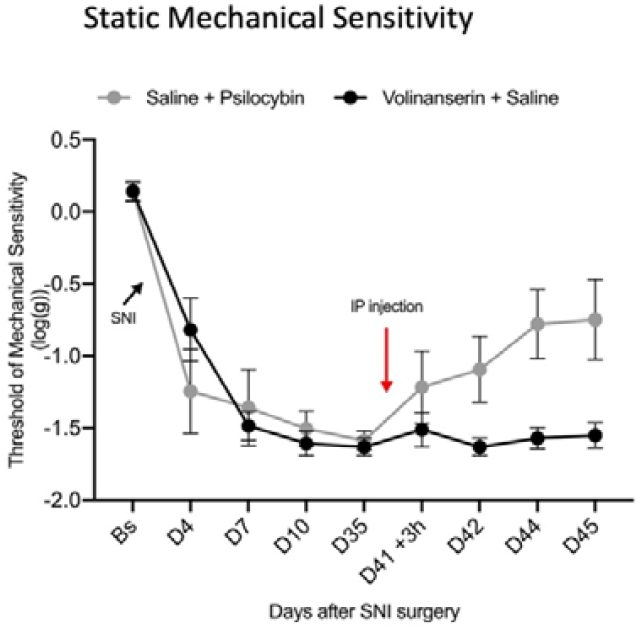
Effect of volinanserin on static mechanical sensitivity. Static mechanical threshold of mice assessed using calibrated von Frey filaments before (BS, baseline) and after SNI surgery. On day 41 after SNI, all mice received an IP injection of saline or volinanserin (0.032mg/kg) and 30 minutes later an injection of psilocybin (0.3 mg/kg) or saline, red arrows (n=5/5).

**Fig S5.**
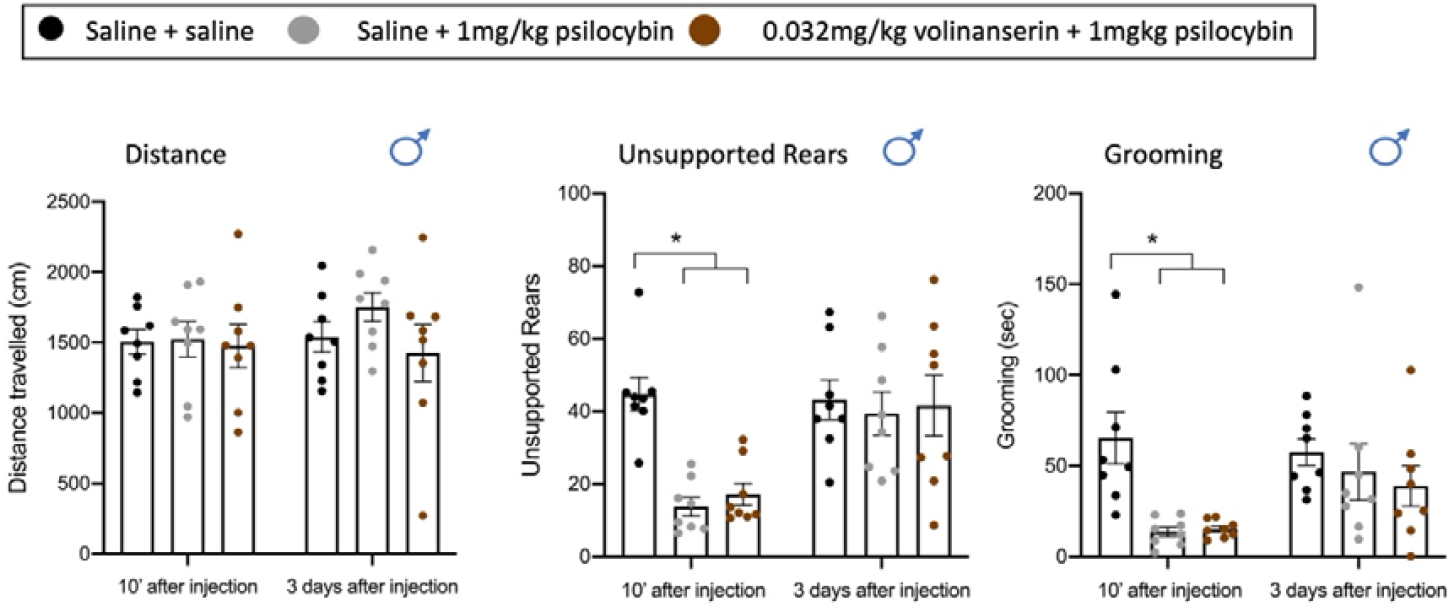
Effects of psilocybin and 5-HT_2A_R blockade on spontaneous behaviours in male mice. Male mice were injected with saline + saline (black), saline + psilocybin (1 mg/kg; grey), or volinanserin (0.032 mg/kg) + psilocybin (1 mg/kg; brown). Spontaneous behaviours were assessed either 10 minutes or 3 days after injection. (i) Total distance travelled (locomotor activity) was not significantly altered by psilocybin or volinanserin at either timepoint. (ii) Unsupported rearing was reduced 10 minutes after psilocybin administration, with or without volinanserin, but returned to baseline by 3 days post-injection. (iii) Grooming duration was similarly reduced 10 minutes after psilocybin and/or volinanserin treatment, with no persistent effects at 3 days. (n=8 pre group, two-way Repeated measured with Mixed Models ANOVA, factor TREATMENT 10 minuets to D3: unsupported rearing, F= 4.5, P=0.023; Tukey HSD sal+sal vs sal/psi p=0.028; sal/sal vs vol/psi p=0,07; grooming, F= 6.6, P=0.006; Tukey HSD sal+sal vs sal/psi p=0.019; sal/sal vs vol/psi p=0,01). Data are presented as mean ± SEM with individual values shown. ^*^p < 0.05.

## References

1. Mills, S. E. E., Nicolson, K. P. & Smith, B. H. Chronic pain: a review of its epidemiology and associated factors in population-based studies. Br J Anaesth 123, e273–e283 (2019).

2. Finnerup, N. B. et al. Pharmacotherapy for neuropathic pain in adults: a systematic review and meta-analysis. The Lancet Neurology 14, 162–173 (2015).

3. Volkow, N. D. & McLellan, A. T. Opioid Abuse in Chronic Pain — Misconceptions and Mitigation Strategies. N Engl J Med 374, 1253–1263 (2016).

4. Robinson, E. S. J. Translational new approaches for investigating mood disorders in rodents and what they may reveal about the underlying neurobiology of major depressive disorder. Phil. Trans. R. Soc. B 373, 20170036 (2018).

5. Bair, M. J., Robinson, R. L., Katon, W. & Kroenke, K. Depression and Pain Comorbidity: A Literature Review. Arch Intern Med 163, 2433 (2003).

6. Hanks, J. B. & González-Maeso, J. Animal models of serotonergic psychedelics. ACS Chem Neurosci 4, 33–42 (2013).

7. Madsen, M. K. et al. Psychedelic effects of psilocybin correlate with serotonin 2A receptor occupancy and plasma psilocin levels. Neuropsychopharmacol. 44, 1328–1334 (2019).

8. Carhart-Harris, R. et al. Trial of Psilocybin versus Escitalopram for Depression. N Engl J Med 384, 1402–1411 (2021).

9. Daws, R. E. et al. Increased global integration in the brain after psilocybin therapy for depression. Nat Med 28, 844–851 (2022).

10. Ramachandran, V., Chunharas, C., Marcus, Z., Furnish, T. & Lin, A. Relief from intractable phantom pain by combining psilocybin and mirror visual-feedback (MVF). Neurocase 24, 105–110 (2018).

11. Schindler, E. A. D. et al. Exploratory Controlled Study of the Migraine-Suppressing Effects of Psilocybin. Neurotherapeutics 18, 534–543 (2021).

12. Kolbman, N. et al. Intravenous psilocybin attenuates mechanical hypersensitivity in a rat model of chronic pain. Current Biology 33, R1282–R1283 (2023).

13. Castellanos, J. P. et al. Chronic pain and psychedelics: a review and proposed mechanism of action. Reg Anesth Pain Med 45, 486–494 (2020).

14. Hashmi, J. A. et al. Shape shifting pain: chronification of back pain shifts brain representation from nociceptive to emotional circuits. Brain 136, 2751–2768 (2013).

15. Askey, T., Lasrado, R., Maiarú, M. & Stephens, G. J. Psilocybin as a novel treatment for chronic pain. British J Pharmacology bph.17420 (2024) doi:10.1111/bph.17420.

16. Decosterd, I. & Woolf, C. J. Spared nerve injury: an animal model of persistent peripheral neuropathic pain. Pain 87, 149–158 (2000).

17. Pertin, M., Gosselin, R.-D. & Decosterd, I. The Spared Nerve Injury Model of Neuropathic Pain. in Pain Research (ed. Luo, Z. D.) vol. 851 205–212 (Humana Press, Totowa, NJ, 2012).

18. Halberstadt, A. L., Chatha, M., Klein, A. K., Wallach, J. & Brandt, S. D. Correlation between the potency of hallucinogens in the mouse head-twitch response assay and their behavioral and subjective effects in other species. Neuropharmacology 167, 107933 (2020).

19. Shao, L.-X. et al. Psilocybin induces rapid and persistent growth of dendritic spines in frontal cortex in vivo. Neuron 109, 2535-2544.e4 (2021).

20. Maiarù, M. et al. Substance P-Botulinum Mediates Long-term Silencing of Pain Pathways that can be Re-instated with a Second Injection of the Construct in Mice. The Journal of Pain S1526590024003407 (2024) doi:10.1016/j.jpain.2024.01.331.

21. Caporoso, J. et al. A Thermal Place Preference Test for Discovery of Neuropathic Pain Drugs. ACS Chem. Neurosci. 11, 1006–1012 (2020).

22. Matsumoto, K. et al. Juvenile social defeat stress exposure favors in later onset of irritable bowel syndrome-like symptoms in male mice. Sci Rep 11, 16276 (2021).

23. Shahar, O. et al. Role of 5-HT2A, 5-HT2C, 5-HT1A and TAAR1 Receptors in the Head Twitch Response Induced by 5-Hydroxytryptophan and Psilocybin: Translational Implications. IJMS 23, 14148 (2022).

24. Haberzettl, R., Fink, H. & Bert, B. Role of 5-HT1A- and 5-HT2A receptors for the murine model of the serotonin syndrome. Journal of Pharmacological and Toxicological Methods 70, 129–133 (2014).

25. Drake, R. A., Steel, K. A., Apps, R., Lumb, B. M. & Pickering, A. E. Loss of cortical control over the descending pain modulatory system determines the development of the neuropathic pain state in rats. eLife 10, e65156 (2021).

26. Baliki, M. N., Mansour, A. R., Baria, A. T. & Apkarian, A. V. Functional Reorganization of the Default Mode Network across Chronic Pain Conditions. PLoS ONE 9, e106133 (2014).

27. Corder, G. et al. An amygdalar neural ensemble that encodes the unpleasantness of pain. Science 363, 276–281 (2019).

28. Goodwin, G. M. et al. Single-Dose Psilocybin for a Treatment-Resistant Episode of Major Depression. N Engl J Med 387, 1637–1648 (2022).

29. Gregory, N. S. et al. Psilocybin has no immediate or persistent analgesic effect in acute and chronic mouse pain models. Preprint at 10.1101/2025.07.06.663398 (2025).

30. Koseli, E. et al. Effect Of Hallucinogenic And Non-Hallucinogenic Psychedelic Analogs On Chronic Neuropathic Pain In Mice. The Journal of Pain 24, 6–7 (2023).

31. Hesselgrave, N., Troppoli, T. A., Wulff, A. B., Cole, A. B. & Thompson, S. M. Harnessing psilocybin: antidepressant-like behavioral and synaptic actions of psilocybin are independent of 5-HT2R activation in mice. Proc. Natl. Acad. Sci. U.S.A. 118, e2022489118 (2021).

32. Husain, M. I. et al. Psilocybin for treatment-resistant depression without psychedelic effects: study protocol for a 4-week, double-blind, proof-of-concept randomised controlled trial. BJPsych open 9, e134 (2023).

33. Vollenweider, F. X., Vollenweider-Scherpenhuyzen, M. F. I., Bäbler, A., Vogel, H. & Hell, D. Psilocybin induces schizophrenia-like psychosis in humans via a serotonin-2 agonist action: NeuroReport 9, 3897– 3902 (1998).

34. Torrado Pacheco, A., Olson, R. J., Garza, G. & Moghaddam, B. Acute psilocybin enhances cognitive flexibility in rats. Neuropsychopharmacology 48, 1011–1020 (2023).

35. Rickli, A., Moning, O. D., Hoener, M. C. & Liechti, M. E. Receptor interaction profiles of novel psychoactive tryptamines compared with classic hallucinogens. European Neuropsychopharmacology 26, 1327–1337 (2016).

36. Brunello, C. A., Cannarozzo, C. & Castrén, E. Rethinking the role of TRKB in the action of antidepressants and psychedelics. Trends in Neurosciences 47, 865–874 (2024).

37. Moliner, R. et al. Psychedelics promote plasticity by directly binding to BDNF receptor TrkB. Nat Neurosci 26, 1032–1041 (2023).

38. Wiffen, P. J. et al. Gabapentin for chronic neuropathic pain in adults. Cochrane Database of Systematic Reviews 2020, (2017).

39. Bonnet, U. & Scherbaum, N. How addictive are gabapentin and pregabalin? A systematic review. European Neuropsychopharmacology 27, 1185–1215 (2017).

40. Calder, A. E. & Hasler, G. Towards an understanding of psychedelic-induced neuroplasticity. Neuropsychopharmacol. 48, 104–112 (2023).

41. Finnerup, N. B., Kuner, R. & Jensen, T. S. Neuropathic Pain: From Mechanisms to Treatment. Physiological Reviews 101, 259–301 (2021).

42. Percie Du Sert, N. et al. The ARRIVE guidelines 2.0: Updated guidelines for reporting animal research. British J Pharmacology 177, 3617–3624 (2020).

43. Chaplan, S. R., Bach, F. W., Pogrel, J. W., Chung, J. M. & Yaksh, T. L. Quantitative assessment of tactile allodynia in the rat paw. J. Neurosci. Methods 53, 55–63 (1994).

44. Duan, B. et al. Identification of Spinal Circuits Transmitting and Gating Mechanical Pain. Cell 159, 1417–1432 (2014).

45. Maiarù, M. et al. The stress regulator FKBP51 drives chronic pain by modulating spinal glucocorticoid signaling. Sci Transl Med 8, 325ra19 (2016).

46. Mills, C. et al. Estimating Efficacy and Drug ED50’s Using von Frey Thresholds: Impact of Weber’s Law and Log Transformation. The Journal of Pain 13, 519–523 (2012).

47. Maiarù, M. et al. Selective neuronal silencing using synthetic botulinum molecules alleviates chronic pain in mice. Sci Transl Med 10, eaar7384 (2018).

